# Synthetic promoter based azide biosensor toolkit to advance chemical-biology

**DOI:** 10.1101/2020.07.08.193060

**Authors:** Chandra Kanth Bandi, Kyle S. Skalenko, Ayushi Agrawal, Neelan Sivaneri, Margaux Thiry, Shishir P.S. Chundawat

**Affiliations:** Department of Chemical & Biochemical Engineering, Rutgers The State University of New Jersey, Piscataway, NJ 08854, USA; Department of Genetics and Waksman Institute, Rutgers The State University of New Jersey, Piscataway, NJ 08854, USA

**Keywords:** Azide biosensor, promoter engineering, protein expression, glycoengineering, chemical-biology

## Abstract

Real-time azide or azido-functionalized molecular detection inside living cells using bioorthogonal chemistry-based approaches has been revolutionary to advancing chemical-biology. These methods have enabled diverse applications ranging from understanding the role of cellular glycosylation pathways, identifying diseased cells, and targeting delivery of azido-based therapeutic drugs. However, while classical techniques were applicable only to *in-vitro* detection of such functional groups, even recent bioorthogonal based-detection methods require expensive sensing reagents and also cannot selectively identify inorganic azide. Here, we report an *in-vivo* synthetic promoter based azide biosensor toolkit to selectively detect azide anions. A promiscuous cyanate-specific promoter was engineered to detect azide and rapidly induce expression of green fluorescent protein (GFP) in *Escherichia coli*. Our synthetic azide operon allows highly-tunable GFP expression, outperforming the classic *lac*-operon, and also offers an alternative low-cost protein expression system. Finally, we showcase the utility of this toolkit for *in-vivo* bioorthogonal reaction biosensing and glycoengineering based applications.

---

Azide ion is an excellent nucleophile that can readily participate in nucleophilic substitution reactions ^1^. This facilitated the discovery and synthesis of organic azides that are ubiquitously used as building blocks in synthetic chemistry and material sciences for over 150 years ^2^. Azides have found widespread commercial applications ranging from automobile airbag propellants, biocides, and as functional groups in fine chemicals and pharmaceuticals ^3,4^. In particular, due to its excellent bactericidal and fungicidal properties, azides have been used as potent antibiotics (e.g., solithromycin, colecoxib, rofecoxib and chloramphenicol) ^5–8^. Many azido containing pharmaceutical drugs such as azidamfenicol, azidocillin, and zidovudine have also been clinically approved for oral use. With the recent invention of biorthogonal click chemistry ^9^, which utilizes alkyne-azide chemistry for diverse applications in chemical biology, drug development, and bioorganic chemistry, there is a reemergence of interest towards organic azides within the scientific community. Azide functionalized probes are actively being developed as molecular-imaging reagents and biomarkers for disease detection ^10^,therapeutic drug delivery ^11^, glycomics/metabolomics ^12^, and chemoproteomics ^13^.

However, the *in vivo* use of azido-based drugs or molecular probes also necessitates the need to monitor organic azide stability for release of toxic azide ions. While organic azides are less toxic, their degradation to release azide ions can lead to acute toxicity. For example, zidovudine drug toxicity was shown to be closely dependent on azido group stability ^14^. The current state-of-the-art *in vitro* techniques involving spectrophotometry ^15,16^, fluorescence ^17–20^, mass spectrometry, ^21,22^ and redox sensing ^23^ have all been used for azide ion detection. However, such *in vitro* detection methods require extensive sample derivatization steps and are not suited for high throughput *in vivo* based detection of azides. There are currently no methods available that do not require additional chemical derivatization for highly selective *in vivo* azide ion detection.

A class of organic azides, glycosyl azides, has facilitated the study of dynamic imaging of carbohydrate-based biomolecules in living systems to understand their function, localization, and pathway regulation ^24^. Furthermore, glycosyl azides are also efficient donor substrates for engineered glycosidases called glycosynthases that can promote glycosidic bond synthesis through transglycosylation for glycans synthesis ^25,26^. Compared to expensive and poorly soluble donor sugar substrates (e.g., glycosyl fluorides and p-nitrophenyl glycosides), glycosyl azides carry a small leaving group with strong nucleophilic character and have good water solubility making them ideal substrates for glycosidase enzymes. However, there are currently limited options available for azide detection as opposed to rapid colorimetric methods for detection of other glycosyl leaving groups (like p-nitrophenyl or pNP) ^27^. This has limited the widespread use of glycosyl azides as reagents for glycosidase activity characterization. Therefore, availability of a synthetic biology toolkit for *in vivo* azide ion detection can broadly enable the fields of chemical glycobiology and glycoengineering.

Here, we have developed an *in vivo* cell based biosensing toolkit to selectively detect azide ions. Briefly, a native *Escherichia coli* cyanate operon was engineered to generate a synthetic promoter plasmid that is selectively inducible by azide ions. The tunable expression of model green fluorescence protein (GFP) was shown using azide based promoter induction and compared to the standard *E. coli* lactose (*lac)* operon. Finally, the biosensing potential of this promoter to selectively detect inorganic azide ions, with respect to organic glycosyl azides, was showcased as a proof-of-concept glycobiology application.

*The E. coli* genome consists of several operons (630-700 operons ^28^) performing specific functions that are essentially for bacterial metabolism and survival in harsh environments. For example, the cyanate or *cyn* operon enables *E. coli* to overcome the toxicity of exogenous cyanate to survive in cyanate-rich environment ^29^. This gene likely evolved in archaea and cyanobacteria for energy production and nitrogen assimilation during the early history of single-celled marine life on earth ^30^. The *cyn* operon in the *E. coli* genome, analogous to the *lac* operon, is comprised of three structural genes; *cynT*, *cynS*, and *cynX* (**Figure 1A**) that encode carbonic anhydrase, cyanate hydratase, and cyanate transporter proteins, respectively. Along with *cynTSX* genes, *cynR* repressor gene encoding cynR protein is present as part of *cyn* operon on the opposite strand. The operon is under tight negative regulation of a cynR repressor protein which upon binding to the operator region results in unfavorable DNA bending to prevent transcription of the *cynTSX* genes. The *cyn* operon is activated when a cyanate molecule binds to the allosteric repressor protein bound to the promoter/operator region. The exogenous cyanate binds to the cynR inducer binding domain and causes conformational changes in the DNA binding domain thereby reducing the bend in the operator region to facilitate transcription ^29,31,32^. The expressed cyanate hydratase catalyzes the bicarbonate-dependent decomposition of cyanate ions into carbon dioxide and ammonia to rapidly decrease the intracellular concentration of cyanate.

**Figure 1.**
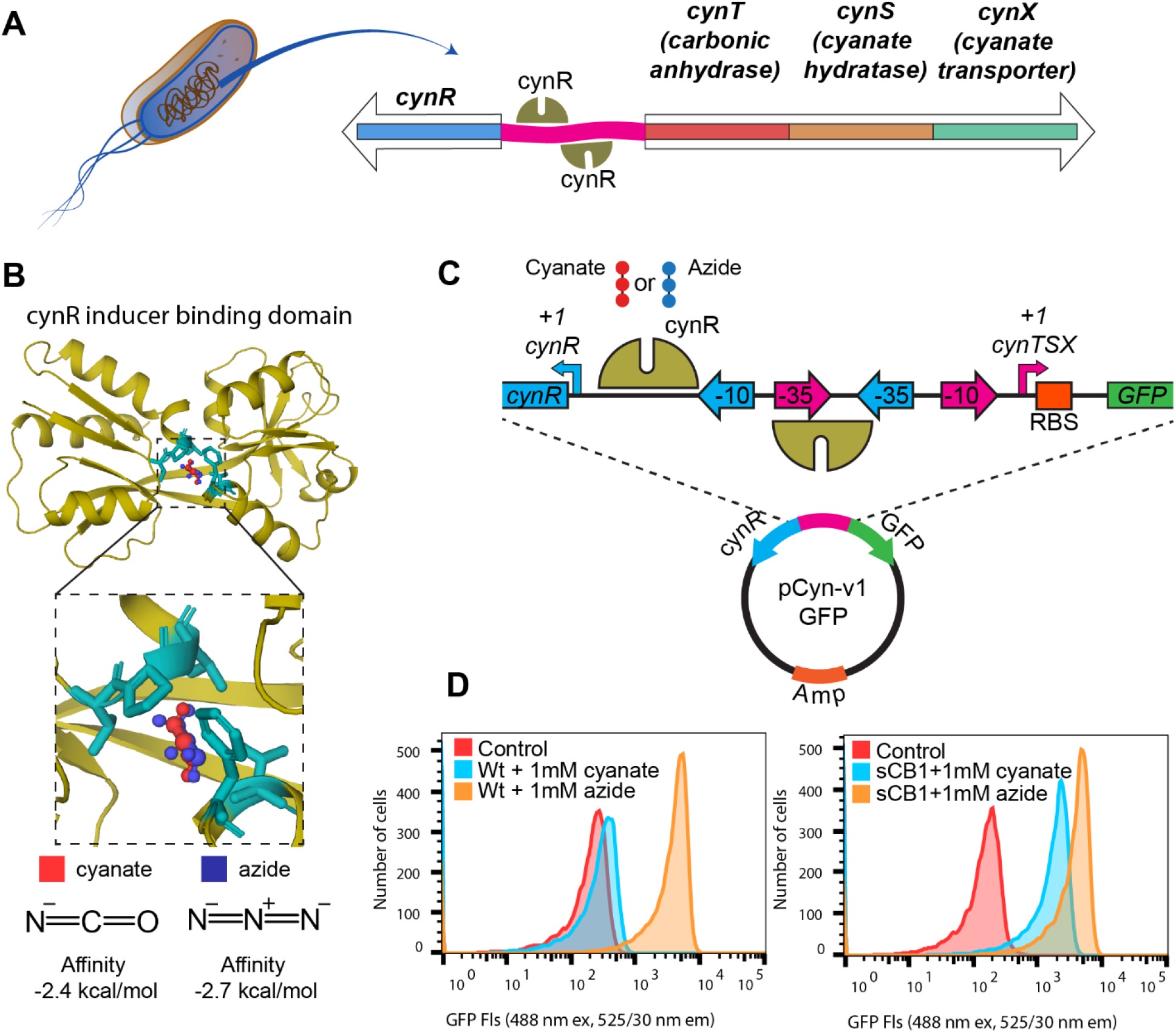
Promoter engineering to enable reporter GFP expression in E. coli using azide and cyanate as inducers. **A.** Native cynTSX operon is present in E. coli genome to facilitate cell growth in a cyanate-rich environment. Cyanate molecule binds to the cynR repressor protein which regulates the downstream protein expression of three essential genes (cynT, cynS, cynX). **B.** Molecular docking of cyanate and azide ions in the binding pocket of cynR protein is shown. Since azide is a structural homolog for cyanate, it also binds tightly in the cynR binding pocket and in the same location as cyanate. **C.** Plasmid map design of pCyn-v1-GFP containing the native cynR and cyn operator region cloned upstream of a GFP reporter gene. The native version (v1) was used first to monitor GFP expression using azide and cyanate as inducers. **D.** Flow cytometer analysis of GFP fluorescence signal confirms cyanate and azide inducible heterologous in vivo protein expression using native (BW25113-wt) and engineered ΔcynS knockout (BW25113-sCB1) E. coli strains.

Cyanate is a small linear molecule that binds strongly inside the binding pocket of the cynR inducer binding domain. Azide ion is structurally homologous to cyanate ion and can therefore theoretically also bind to CynR. To confirm if azide can bind with similar binding affinity to cynR, Autodock vina ^33^ was used to perform docking simulations for both cyanate and azide in the binding pocket of cynR. The binding orientation of the docked ligands at minimum free energy reveals a good overlap in the binding site with similarly predicted binding affinities (**Figure 1B**). This suggested that azide could also function as a gratuitous inducer for the *cyn* operon. Next, we subcloned the regulatory segment consisting of *cynR* gene and the *cyn* operator region into a plasmid vector with ampicillin resistance and with green fluorescent protein (*gfp*) as a reporter gene. The regulatory segment was placed upstream of the *gfp* reporter gene to facilitate GFP expression using both cyanate and azide as inducers (**Figure 1C**). The resultant plasmid (called pCyn-v1-GFP) was transformed into *E. coli* BW25113 wildtype (Wt) strain and induced with 1 mM each of sodium cyanate or sodium azide. The supernatant of lysed cells after induction were analyzed for fluorescence using a spectrophotometer (details available in supporting information or **SI Figure S1**). In addition, the fluorescence of individual intact cells were analyzed through a flow cytometer and **Figure 1D** presents the resultant histogram plots of the induced cells fluorescence after 19 hours. Compared to the control (uninduced cells), the azide induced cell lysate had a nearly 20-fold increase in the fluorescence while cyanate did not result in any significant GFP expression. The absence of GFP expression in cyanate induced cells was likely due to the breakdown of cyanate by the endogenous cyanate hydratase enzyme encoded by the native *E. coli* BW25113-Wt cells. Hence, we next tested the induction capacity of the synthetic promoter using a Δ*cynS* knockout strain (BW25113-sCB1) which clearly recovered GFP expression during induction by cyanate (**SI Figure S1**). However, the amount of protein expressed based on total cell lysate GFP fluorescence was very low suggesting that the native *cyn* promoter strength was quite poor. This is not surprising since the native *cyn* promoter regulates associated CynTSX proteins expression required to overcome cyanate toxicity and that seldom requires high protein yields. The native operator region also contains suboptimal −10 sequence and Shine-Delgarno (SD) sequences (or ribosomal binding site; RBS) which could also play a significant role in poor expression strength.

To improve the native promoter strength for enabling inducible higher protein expression levels, the native *cyn* promoter was next engineered to contain consensus −10 and SD sequences. The *cynR* gene, that is negatively regulated by *cyn* promoter, was placed under control of an independent constitutive promoter ^34^(see SI for sequence information). Additional to these modifications, to avoid interference and steric hinderances between the two promoters, the promoters were separated by inserting a random DNA spacer sequence of 100 bp (pCyn-v2-GFP) and 1000 bp (pCyn-v4-GFP) (**Figure 2B** and **SI Figure S2**). The control construct that contains no spacer sequence (pCyn-v3-GFP) was also generated. The modified constructs were individually transformed into *E. coli* cells and tested for GFP expression using sodium azide as inducer. All three engineered constructs showed a 30-fold increase in the fluorescence signal in the cell lysate, indicating a significant increase in GFP expression as compared to the native pCyn-v1-GFP promoter (**Figure 2D** and **SI Figure S2B**). The pCyn-v4-GFP construct with longest spacer region showed around 30% increased fluorescence when compared to the no spacer pCyn-v3-GFP. However, significant leaky GFP expression in the v4 construct was seen for the no induction control (**SI Figure S2C**). The extra-long spacer sequences could be potentially forming undesirable interactions in the plasmid causing a change in the DNA bending properties for CynR.

**Figure 2.**
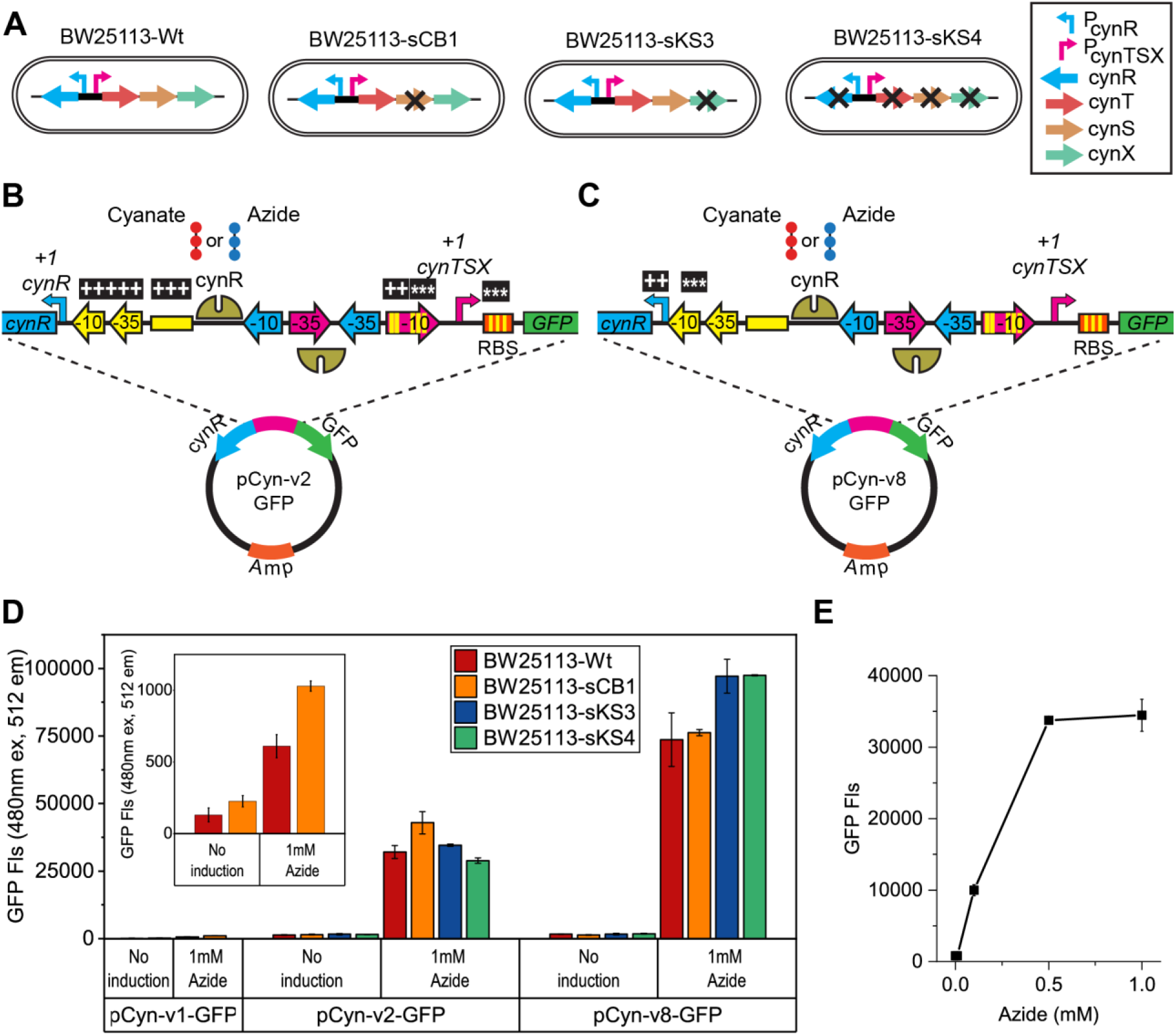
Strain engineering and promoter design co-optimization to increase reporter GFP protein expression upon induction by azide. **A**. E. coli BW25113 wild type and various knockout strains are represented here in the cartoon schematics. P_cynR_ and P_cynTSX_ are promoters for cynR and cynTSX genes, respectively. Individual genes are color coded and knocked-out genes are illustrated with “X” mark. **B&C**. Plasmid map design of pCyn-v2-GFP and pCyn-v8-GFP containing the engineered regulatory region between cynR and GFP genes is highlighted here. The mutational replacements in the promoter region are represented by asterisk (*) while additions are denoted by plus sign (+). **D**. Bar graphs representing GFP fluorescence intensities from cell lysates of engineered strains containing three distinct plasmid variants (v1, v2, v8) is shown. The magnified inset shows GFP fluorescence for native promoter construct (pCyn-v1-GFP). **E.** Fluorescence plot for cell lysate of BW25113-wt strain containing pCyn-v2-GFP and induced with increasing concentrations of sodium azide. Error bars indicate one standard deviation from reported mean values from three biological replicates.

Decreasing the length between the promoters largely reduced the leaky expression under no induction while maintaining the largely improved GFP yield upon azide induction (**Figure S2**). For all the future work, pCyn-v2-GFP design was chosen to keep the promoters at optimal distance from each other. The 30-fold increase in GFP fluorescence observed was now promising to be able to utilize the pCyn-v2-GFP engineered design for sensing azide inside cells. To determine the minimal azide amount required to induce GFP protein expression, we induced cells transformed with the pCyn-v2-GFP plasmid with varying concentrations of azide. The maximum amount of azide used for induction should be limited to 5 mM with the current *E. coli* strains tested since higher amounts show dramatically negative influence on cell growth (**Figure S3**). GFP fluorescence of cell lysate showed a strong correlation in protein expression as a function of inducer dosage. GFP expression increased until 1 mM azide concentration induction after which there was reduction in the GFP signal for 5 mM inducer concentration likely due to cellular toxicity (**Figure 2E**). The lowest tested azide concentration (10 μM) showed very marginal GFP expression indicating that the detection limit for this synthetic promoter would be around 10 μM. The bacterial growth phase during the time of induction also plays an important role and we observed that induction at early exponential phase can yield higher protein amounts (**Figure S4**).

CynR protein acts as a repressor for the promoter by causing a bend at −35 site hindering the binding of RNA polymerase. The cynR protein consists of two domains: an inducer binding domain and a DNA binding domain. The inducer binding domain binds to the inducer (cyanate or azide) which decreases the bend at the promoter site to facilitate transcription. In regulating protein expression, along with the inducer amount, the relative amounts of cynR protein is also critical. We therefore further engineered the cynR constitutive promoter to adjust the background level of repressor protein expressed. Four different constructs (v5, v6, v7 and v8) were constructed by introducing mutations at the promoter −10 site, regions between −10 and − 35 site, and between RBS and translation start site (**Figure 2C** and **SI Figure S5A**). The expression strength of these modified promoters were tested in wild type (BW2113) and additional knockout strains (sCB1, sKS3, sKS4) using GFP fluorescence in cell lysate. The knockout strains sKS3 and sKS4 were generated to remove the *cynX* gene and *cyn* operon respectively (**SI Table S1**) to minimize the export of azide ions using transporter protein and reduce interferences from endogenous *cyn* operon. The pCyn-v8-GFP construct, which had reduced efficiency at −10 site but optimal length between RBS and translation start site, gave about 120-160 fold more in reporter GFP fluorescence as compared to the native promoter, with the maximum fold increase observed in ΔcynX (sKS3), ΔcynR, and ΔcynTSX (sKS4) knockout strains (**Figure 2D**). On the other hand, while the designs (v5, v6, v7) not only did not show any significant increase in fluorescence with respect to v2 design they resulted in leaky background expression without any addition of inducer (**SI Figure S5**).

The optimal plasmid designs (pCyn-v2-GFP and pCyn-v8-GFP) can function as tunable synthetic biosensors for rapid *in vivo* detection of azide. The amount of GFP expressed within 2 hours of 1 mM azide induction in BW25113-Wt strain was estimated to be 0.4 mg and 0.9 mg from 1 ml culture for pCyn-v2-GFP and pCyn-v8-GFP, respectively, based on the calibration curve between protein concentration and GFP fluorescence (**Figure S6**). The protein expression efficiency of the engineered promoter (pCyn-v2-GFP) was next compared against the engineered lac expression system ^35,36^. The synthetic *cyn* promoter in BW25113-Wt (**Figure 3A**) and *lac* promoter in BL21 cells (**Figure 3B**) were induced using sodium azide and IPTG, respectively, at various concentrations ranging between 0.01 mM to 1 mM. The amount of protein expressed in P_lac_ system was higher during the early time points after induction and at the lowest inducer concentrations. Even at lower IPTG concentrations (0.01 mM and 0.1 mM), GFP expression increased with induction time and reached a maximum after 24 hours of induction. Whereas we see a relatively lower (or no) GFP expression at similar low azide concentrations (**Figure 3A** and **3B**). However, we see a tunable expression using azide induction as amount of GFP expressed is closely proportional to inducer concentration, unlike IPTG. Also, the maximum GFP expression achieved after 24 hours was much higher for the *cyn* promoter than *lac* promoter at slightly higher inducer concentrations (>0.5 mM) (**Figure 3A** and **3B**). These results showcase the utility of the engineered plasmid harboring the optimized P_cyn_ promoter as an alternative system for heterologous protein expression.

**Figure 3.**
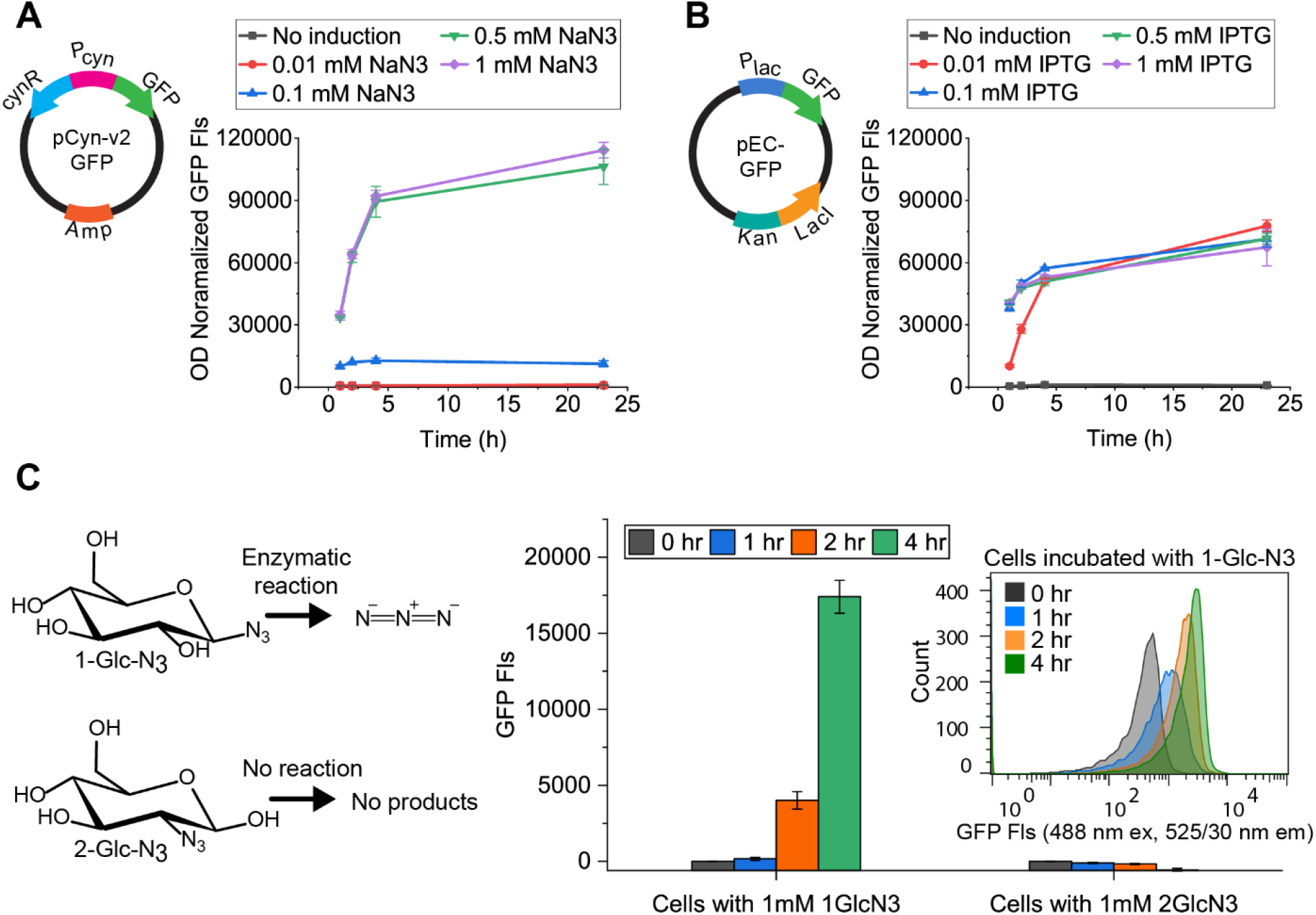
Application of synthetic azide promoter towards heterologous protein expression and biorthogonal chemical-biology. **A**. E. coli BW25113-Wt containing pCyn-v2-GFP plasmid was induced with varying azide concentrations for 24 hours and GFP fluorescence of cell lysate was measured at various time points. **B**. E. coli BL21 containing pEC-GFP plasmid was induced with different IPTG concentrations for 24 hours and GFP fluorescence of the cell lysate was measured at different time points. **C**. E. coli BW25113-Wt cells with pCyn-v2-GFP plasmid was incubated with 1-azido-β-D-glucopyranosyl azide (1-Glc-N_3_) and 2-Deoxy-2-azido-β-D-glucopyranosyl azide (2-Glc-N_3_) and GFP fluorescence of respective cell lysates are shown as bar graphs. The inset depicts the flow cytometry analysis data for cells incubated with 1-Glc-N3 for 4 hours. Error bar indicates one standard deviation from reported mean values from three biological replicates.

In addition to application as a heterologous expression system, the developed azide promoter can be useful for chemical biologists especially in glycobiology and glycosynthase engineering related applications. This is a powerful synthetic biology tool that will allow selective identification of inorganic azides amongst other chemically linked organic azides. Enzymatic reactions which result in the release of azide ion can be potentially identified using this promoter system. Carbohydrate-Active enZymes (CAZymes) such as glycosyl hydrolases and transglycosidases have been shown to release azide from azido-based sugars during glycosidic bond hydrolysis ^37^ and synthesis ^26,37–40^, respectively. Multiple glycosyl azides such as galactosyl-, glucosyl-, and mannosyl-azides have been reported to be hydrolyzed by galactosidases, glucosidases, and mannosidases, respectively ^37^. Similarly, engineered glycosidases (i.e., transglycosidases and glycosynthases) have been shown to use azido-hexoses and their N-acetyl derivatives as donor sugars for oligosaccharides synthesis ^38,40^. β-glucosidases from *Aspergillus* sp. and *Agrobacterium* sp. ^39^ belonging to glycosyl hydrolase families GH3 and GH1 were reported to actively cleave azide group from glucosyl azide. Similarly, homologous β-glucosidases from the GH1 and GH3 families present in *E. coli* can potentially also show similar substrate specificity to release azide when incubated with glucosyl azide. Here, we incubated *E. coli* BW25113-Wt cells containing the engineered pCyn-v2-GFP plasmid with 1-azido-β-D-glucopyranosyl azide (1-Glc-N3) for 4 hours. As illustrated in **Figure 3C**, a small change in GFP fluorescence was seen in the first hour after which a rapid increase in fluorescence was observed owing to the release of azide when using 1-Glc-N3. To verify if the detected GFP fluorescence is due to cleavage of azide and not due to presence of azido-glucose, we incubated the cells with 2-Deoxy-2-azido-β-D-glucopyranosyl azide (2-Glc-N3) analogous to the 2-deoxy-2-fluoro glycoside based β-glucosidase inhibitors ^41^. No change in fluorescence was detected in the control samples since 2-Glc-N3 is not hydrolyzed by β-glucosidases and hence cannot release free azide for induction of the promoter. The preferential sensitivity of the designed promoter towards azide ions can be used for *in vivo* engineering of efficient CAZymes for sugar polymer synthesis and hydrolysis ^27^.

In summary, we have demonstrated the design and engineering of an azide inducible promoter system for *E. coli*. This azide promoter system allows tunable expression by varying the inducer (azide ion) concentration and outperforms conventional lactose/IPTG based system for heterologous reporter GFP expression. Additionally, the developed toolkit functions as a biosensor for detecting the presence of azide ions inside living cells. This would allow employing this tool for engineering CAZymes such as glycosyl hydrolases which use azido-sugars as substrates and autonomously monitor released azide ions upon substrate hydrolysis or transglycosylation. Furthermore, this biosensing system can be evolved for use with other prokaryotic or eukaryotic cell protein expression systems by replacing the current −10 and −35 sequences with specific target RNA polymerase recognition sequence. Adaptation of an azide specific promoter system for other cell types would be beneficial overall for diverse drug development and chemical-biology focused research communities.

## References

1. Soli, E. D. et al. Azide and cyanide displacements via hypervalent silicate intermediates. J. Org. Chem. 64, 3171–3177 (1999).

2. Brse, S. & Banert, K. Organic Azides: Syntheses and Applications. Technology (John Wiley & Sons, Ltd, 2009). doi:10.1002/9780470682517

3. Madlung, A. The Chemistry behind the Air Bag: High Tech in First-Year Chemistry. J. Chem. Educ. 73, 347 (1996).

4. Dyukarev, E. A. & Knyazeva, A. G. Model of detonation of lead azide (Pb(N3)2) with regard to fracture. Int. J. Fract. 100, 197–205 (1999).

5. Wereide, K. Sensitivity to azidamphenicol. Contact Dermatitis 1, 271–272 (1975).

6. Bengtsson, E., Strandell, T., Svanbom, M. & Tunevall, G. Azidocillin Treatment of Enterococcal Septicemia. Scand. J. Infect. Dis. 4, 143–148 (1972).

7. Blum, M. R., Liao, S. H., Good, S. S. & de Miranda, P. Pharmacokinetics and bioavailability of zidovudine in humans. Am. J. Med. 85, 189–94 (1988).

8. Glassford, I. et al. Ribosome-Templated Azide-Alkyne Cycloadditions: Synthesis of Potent Macrolide Antibiotics by in Situ Click Chemistry. J. Am. Chem. Soc. 138, 3136–3144 (2016).

9. Devaraj, N. K. The Future of Bioorthogonal Chemistry. ACS Cent. Sci. 4, 952–959 (2018).

10. Devaraj, N. K., Thurber, G. M., Keliher, E. J., Marinelli, B. & Weissleder, R. Reactive polymer enables efficient in vivo bioorthogonal chemistry. Proc. Natl. Acad. Sci. 109, 4762–4767 (2012).

11. Au, K. M., Wang, A. Z. & Park, S. I. Pretargeted delivery of PI3K/mTOR small-molecule inhibitor–loaded nanoparticles for treatment of non-Hodgkin’s lymphoma. Sci. Adv. 6, eaaz9798 (2020).

12. Laughlin, S. T. & Bertozzi, C. R. Imaging the glycome. Proc. Natl. Acad. Sci. 106, 12 LP – 17 (2009).

13. Medina-Cleghorn, D. & Nomura, D. K. Exploring Metabolic Pathways and Regulation through Functional Chemoproteomic and Metabolomic Platforms. Chem. Biol. 21, 1171–1184 (2014).

14. Dunge, A., Chakraborti, A. K. & Singh, S. Mechanistic explanation to the variable degradation behaviour of stavudine and zidovudine under hydrolytic, oxidative and photolytic conditions. J. Pharm. Biomed. Anal. 35, 965–970 (2004).

15. Adhikari, S., Guria, S., Ghosh, A., Pal, A. & Das, D. A curcumin derived probe for colorimetric detection of azide ions in water. New J. Chem. 41, 15368–15372 (2017).

16. Tsuge, K., Kataoka, M. & Seto, Y. Rapid determination of cyanide and azide in beverages by microdiffusion spectrophotometric method. J. Anal. Toxicol. 25, 228–236 (2001).

17. Sahana, A. et al. Highly selective organic fluorescent probe for azide ion: formation of a “molecular ring”. Analyst 137, 1544 (2012).

18. Dhara, K. et al. A new water-soluble copper(II) complex as a selective fluorescent sensor for azide ion. Chem. Commun. 46, 1754–1756 (2010).

19. Wang, K. et al. A metal-free turn-on fluorescent probe for the fast and sensitive detection of inorganic azides. Bioorg. Med. Chem. Lett. 26, 1651–1654 (2016).

20. Puthiyedath, T. & Bahulayan, D. A click-generated triazole tethered oxazolone-pyrimidinone dyad: A highly selective colorimetric and ratiometric FRET based fluorescent probe for sensing azide ions. Sensors Actuators, B Chem. 239, 1076–1086 (2017).

21. Wang, L., Dai, C., Chen, W., Liu Wang, S. & Wang, B. Facile derivatization of azide ions using click chemistry for their sensitive detection with LC-MSwz. Chem. Commun. Chem. Commun 47, 10377–10379 (2011).

22. Dillen, L. et al. Quantitative LC-MS/MS analysis of azide and azidoalanine in in vitro samples following derivatisation with dansyl chloride. Anal. Methods 5, 3136–3141 (2013).

23. Lim, J. Y. C. & Beer, P. D. A Halogen Bonding 1,3-Disubstituted Ferrocene Receptor for Recognition and Redox Sensing of Azide. Eur. J. Inorg. Chem. 2017, 220–224 (2017).

24. Baskin, J. M. et al. Copper-free click chemistry for dynamic in vivo imaging. Proc. Natl. Acad. Sci. U. S. A. 104, 16793–7 (2007).

25. Bojarová, P. & Kren, V. Azido leaving group in enzymatic synthesis-small and efficient. in 168–175 (2010). doi:10.1039/9781849730891-00168

26. Fialová, P. et al. Glycosyl azide—a novel substrate for enzymatic transglycosylations. Tetrahedron Lett. 46, 8715–8718 (2005).

27. Agrawal, A. et al. Click-chemistry enabled directed evolution of glycosynthases for bespoke glycans synthesis. bioRxiv 2020.03.23.001982 (2020). doi:10.1101/2020.03.23.001982

28. Salgado, H., Moreno-Hagelsieb, G., Smith, T. F. & Collado-Vides, J. Operons in Escherichia coli: Genomic analyses and predictions. Proc. Natl. Acad. Sci. U. S. A. 97, 6652–6657 (2000).

29. Sung, Y.-C. & Fuchs, J. A. Characterization of the cyn Operon in Escherichia coli K12* Young-chul Sung and. J. Biol. Chem. 263, 14769–14775 (1988).

30. Palatinszky, M. et al. Cyanate as an energy source for nitrifiers. Nature 524, 105–108 (2015).

31. Sung, Y.-C. & Fuchs, J. A. The Escherichia coli K-12 cyn Operon Is Positively Regulated by a Member of the lysR Family. 174, (1992).

32. Lamblin, A.-F. J. & Fuchs, J. A. Functional Analysis of the Escherichia coli K-12 cyn Operon Transcriptional Regulation. J. BACrERIOLOGY 176, 6613–6622 (1994).

33. Trott,O., Olson, A. J. Autodock vina: improving the speed and accuracy of docking. J. Comput. Chem. 31, 455–461 (2019).

34. Stringer, A. M. et al. FRUIT, a Scar-Free System for Targeted Chromosomal Mutagenesis, Epitope Tagging, and Promoter Replacement in Escherichia coli and Salmonella enterica. PLoS One 7, (2012).

35. Studier, F. W. & Moffatt, B. A. Use of bacteriophage T7 RNA polymerase to direct selective high-level expression of cloned genes. J. Mol. Biol. 189, 113–130 (1986).

36. Eames, M. & Kortemme, T. Cost-Benefit Tradeoffs in Engineered lac Operons. Science (80-.). 336, 911–915 (2012).

37. Bojarová, P. et al. Glycosyl azides - An alternative way to disaccharides. Adv. Synth. Catal. 349, 1514–1520 (2007).

38. Cobucci-Ponzano, B. et al. beta-Glycosyl azides as substrates for alpha-glycosynthases: preparation of efficient alpha-L-fucosynthases. Chem. Biol. 16, 1097–108 (2009).

39. Müllegger, J., Jahn, M., Chen, H. M., Warren, R. A. J. & Withers, S. G. Engineering of a thioglycoligase: Randomized mutagenesis of the acid-base residue leads to the identification of improved catalysts. Protein Eng. Des. Sel. 18, 33–40 (2005).

40. Cobucci-Ponzano, B. et al. A novel -d-galactosynthase from Thermotoga maritima converts -d-galactopyranosyl azide to -galacto-oligosaccharides. Glycobiology 21, 448–456 (2011).

41. Rempel, B. P. & Withers, S. G. Covalent inhibitors of glycosidases and their applications in biochemistry and biology. Glycobiology 18, 570–586 (2008).

