## Supplementary Information for "Synthetic promoter based azide biosensor toolkit to advance chemical-biology"

### for

**This PDF file includes:**

Supplementary Text (including Materials and Methods)
Figures S1 to S6
Tables S1 and S2
SI References

**Materials and Methods:**

**Bacterial strain engineering:**

All the chemicals, reagents, and solvents were purchased from Fisher Scientific and Sigma-Aldrich and used without purification. *E. coli* strain used for cloning was E. cloni® 10G (Lucigen, WI, USA) and for protein expression using lac promoter was BL21-CodonPlus-RIPL [λDE3] (Stratagene, Santa Clara, CA, USA). Phusion 2X high fidelity PCR master mix (0.04 U/µL Phusion DNA polymerase, 400 µM dNTPs, Phusion 2XHF buffer, 3 mM MgCl2) was purchased from Thermo Fisher Scientific (USA) and restriction enzymes were procured from New England BioLabs Inc. (USA). The primers used for site directed mutagenesis (SDM), sequence and ligation independent cloning (SLIC), and sequencing reactions were obtained from Integrated DNA Technologies, Inc (USA). Successfully cloned plasmids were isolated from E. cloni® 10g cells using IBI Scientific (USA) plasmid extraction kit and sequences were confirmed through Sanger sequencing performed by Genscript Inc. (NJ, USA). The carbohydrate substrates used in reported assays were purchased from Synthose Inc, Canada.

The *E.coli* strains used for protein expression using *cyn* promoter were constructed using the protocol and reagents outlined in Datsenko 2000 [1]. Briefly, strains sCB1 (BW25113 *cynS*::FRT) and sKS3 (BW25113 *cynX*::FRT) were constructed by first streaking out the strains JW0331 (BW25113 *cynS*::Kan) and JW0332 (BW25113 *cynX*::Kan)[2] respectively from the Keio collection onto LB-agar kanamycin (50 µg/ml) plate. The kanamycin marker was then removed by transforming the individual strains with pCP20, following the protocol outlined in Datsenko 2000, and curing the strain of the plasmid. The final strain was diagnosed by PCR and for loss of kanamycin resistance.

Strain sKS4, BW25113 *cynR,cynTSX*::FRT, was constructed by first transforming the Keio parent strain, BW25113, with pKD46 and plated onto carbenicillin (100 µg/mL) plates. 5 mL overnights of BW25113 + pKD46 in LB + carbenicillin (100 µg/mL) were grown at 30°C and then back-diluted 1:100 into 50 mL LB + carbenicillin (100 µg/mL) + 0.2% L-arabinose. The 50 mL culture was grown to an OD600 of 0.8, at 30°C, and cells were washed 4 times with 50 mL of ice-cold water. The final cell pellet was resuspended 1:250$^{th}$ the starting culture volume with fresh water and sat on ice until ready to electroporate. The linear DNA fragment, that was used to knockout the *cynR* gene and *cynTSX* operon, was synthesized by amplifying the kanamycin gene from pKD4 using primers k1 and k2. The linear DNA fragment was checked by gel electrophoresis for purity and was then cleaned-up and concentrated using Qiagen's PCR clean-up kit, the product was eluted in water. The intermediate strain sKS1 (BW25113 *cynR*, *cynTSX*::Kan) was made by mixing 100 ng of linear DNA with 50 µL electrocompetent BW25113 + pKD46 cells, shocking immediately with 1.8 kV (with a pulse constant of 5.2ms), and recovered in 900 µL of SOC at 37°C for 3 hours. After recovery, cells were spun down at 10,000xg for 2 minutes at room temperature, resuspended to 100 µL, plating solution onto kanamycin (50 µg/mL) plates, and grown at 37°C overnight. sKS1 was transformed with pCP20 to remove the kanamycin selection marker, following the protocol in Datsenko 2000, making the final strain sKS4 (BW25113 *cynR,cynSTX*::FRT). sKS4 was diagnosed by loss of kanamycin resistance and PCR.

**Design and construction of pCyn vectors:**

The cloning of the plasmid constructs used in this study were performed using Sequence and Ligation-Independent Cloning (SLIC) protocol as outlined in Stevenson et.al [3]. Briefly, to create the pCyn-v1-GFP, the gene fragment consisting of native cynR and cyn promoter/operator region (gblock1) was custom synthesized from Genscript Inc, USA. The GFP gene fragment was taken from pEC-GFP plasmid available at Chundawat lab. Both the gene fragments were cloned into the parent plasmid, ptrc99a while getting rid of the intrinsic lac promoter using the primers p1-p4 and following the SLIC protocol (**SI Table S2A**). To generate pCyn-v2-GFP, an optimized promoter region was designed in-silico and the corresponding DNA fragment (gblock2) was custom synthesized and pCynv-1-GFP was used as starting DNA with primers used being p5-p8. The spacer region of 100 bp from the pCyn-v2-GFP was removed using primers p9-p10 to get pCyn-v3-GFP while an additional random sequence of 900 bp (gblock3) was added using p11-p14 to generate pCyn-v4-GFP. Further, pCyn-v5-GFP was constructed from pCyn-v2-GFP by site directed mutagenesis of the cynR constitutive promoter using primers p15-p16. In the end, the pCyn-v5-GFP was the parent DNA used to create pCyn-v6-GFP, pCyn-v7-GFP and pCyn-v8-GFP using the primers p17-p18, p19-p20 and p21-p22, respectively. All the constructs were diagnosed using Sanger sequencing and the sequence verified plasmids were preserved at -80°C and their corresponding transformed E. cloni cells were stored in 15% glycerol stocks at -80°C.

**Induction of pCyn-v1/v2-GFP expression:**

The pCyn-GFP plasmid constructs were transformed into BW25113 strains (wt, sCB1) and individual colonies were obtained on LB agar plates supplemented with 100 µg/ml Carbenicillin antibiotic. The transformants were inoculated into 10 ml of LB media with carbenicillin and grown at 37°C for 16 hrs. The 10 ml of overnight grown culture was transferred to 200 ml of fresh LB media with carbenicillin and incubated at 37°C until mid-exponential phase (OD 0.4-0.6) was reached. At this point, the 200 ml culture was split into six 25 ml falcon tubes. Two of the tubes were induced with 1 mM sodium azide and two tubes with 1 mM cyanate while the remaining two tubes were used as no induction control. The cultures were placed in the 37°C shaking incubator and 2 ml of sample for bulk fluorescence measurement and 200 ul of sample for flow cytometer runs were collected at every time point.

**Induction of pCyn-v2/v3/v4-GFP expression:**

The pCyn-GFP plasmid constructs were transformed into BW25113 strains (wt, sCB1, sKS3, sKS4) and individual colonies were obtained on LB agar plates supplemented with 100 µg/ml Carbenicillin antibiotic. The transformants were inoculated into 10 ml of LB media with carbenicillin and grown at 37°C for 16 hrs. The 10 ml of overnight grown culture was transferred to 200 ml of fresh LB media with carbenicillin and incubated at 37°C until mid-exponential phase (OD 0.4-0.6) was reached. At this point, the 200 ml culture was split into six 25 ml falcon tubes. Three of the tubes were induced with 1 mM sodium azide and remaining three tubes were used as no induction control. The cultures were placed in the 37°C shaking incubator and 2 ml of sample was collected at every time point for bulk fluorescence measurement and 200 ul was sampled to measure OD600.

**Induction of pCyn-v2/v5/v6/v7/v8-GFP expression:**

The pCyn-GFP plasmid constructs were transformed into BW25113 strains (wt, sCB1, sKS3, sKS4) and individual colonies were obtained on LB agar plates supplemented with 100 µg/ml Carbenicillin antibiotic. The transformants were inoculated into 10 ml of LB media with carbenicillin and grown at 37°C for 16 hrs. Then, 400 µl of overnight grown culture was transferred to 20 ml of fresh LB media with carbenicillin and incubated at 37°C until an OD600 of 0.3-0.4 was reached. At this point, 1 ml of culture was added to 12 wells in a 96 deep well plate. Three wells each were induced with 10 µM, 100 µM 1000 µM of sodium azide and three wells were left uninduced. The

96 well plate was placed in the 37°C shaking incubator for 2 hours. 200 ul of the sample was used for OD600 measurements and the residual sample was used for bulk GFP fluorescence measurement.

**Bulk GFP fluorescence measurement:**

The samples collected from each experiment were first centrifuged to remove the LB media. The pelleted cells were resuspended in 250 µl of BPER reagent (Bacterial Protein Extraction Reagent) and incubated at room temperature for 10 min to lyse the cells. The resultant samples were centrifuged and 200 ul of the supernatant was used to measure GFP fluorescence in a black opaque bottom 96 well plate using a spectrophotometer.

**Flow cytometer data acquisition and analysis:**

For measuring the GFP expression in individual cells, flow cytometer analysis was performed as reported previously by the Chundawat lab [4]. Briefly, at each timepoint for a given induction experiment, 200 µl of cells were collected. The cells were centrifuged to remove the LB media components, resuspended in 1x PBS buffer (phosphate buffered saline) and run through the Guava® easyCyte™ flow cytometer to measure the fluorescence distribution at 488 nm excitation and 525 nm emission. Guavasoft 3.3 software was used for gating live cells based on forward scatter (FSC) and side scatter (SSC) and the fluorescence associated with each cells was collected for 10,000 cells per sample and analyzed using FlowJo software.

**Supplementary Figure S1.** GFP fluorescence of cell lysate measured using spectrophotometer after inducing BW25113-Wt and BW25113-sCB1 strains containing pCyn-v1-GFP with 1 mM cyanate and azide. Error bars indicate one standard deviation from reported mean values from two biological replicates

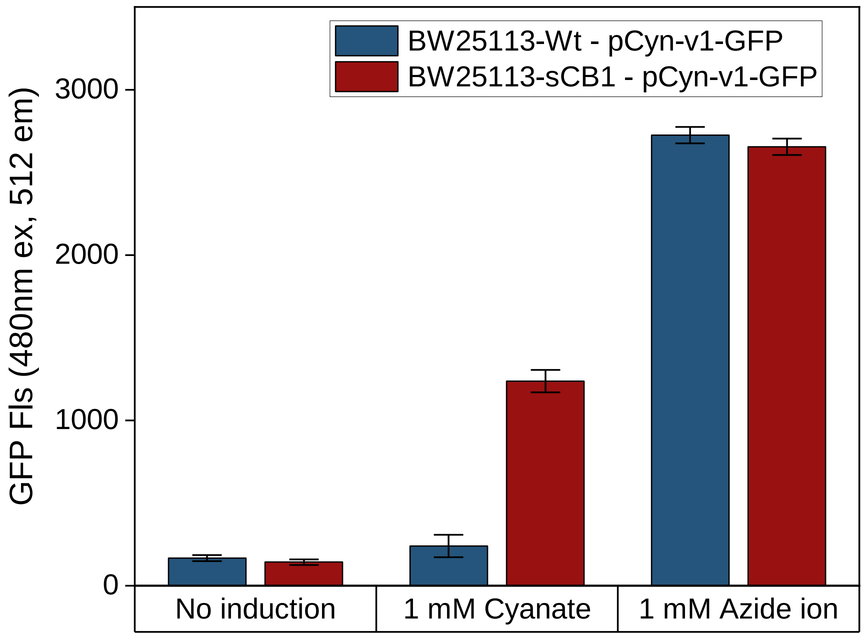

**Supplementary Figure S2. A.** Illustration of the promoter regions of pCyn-v1-GFP, pCyn-v2-GFP, pCyn-v3-GFP, pCyn-v4-GFP. Fluorescence measurements of lysates from BW25113-Wt cells containing plasmids pCynv2-GFP (blue square), pCyn-v3-GFP (red triangle), and pCyn-v4-GFP (green circle) and induced with **B.** 1 mM sodium azide, or **C.** uninduced cells for 4 hours at 37°C. Error bars indicate one standard deviation from reported mean values from three biological replicates

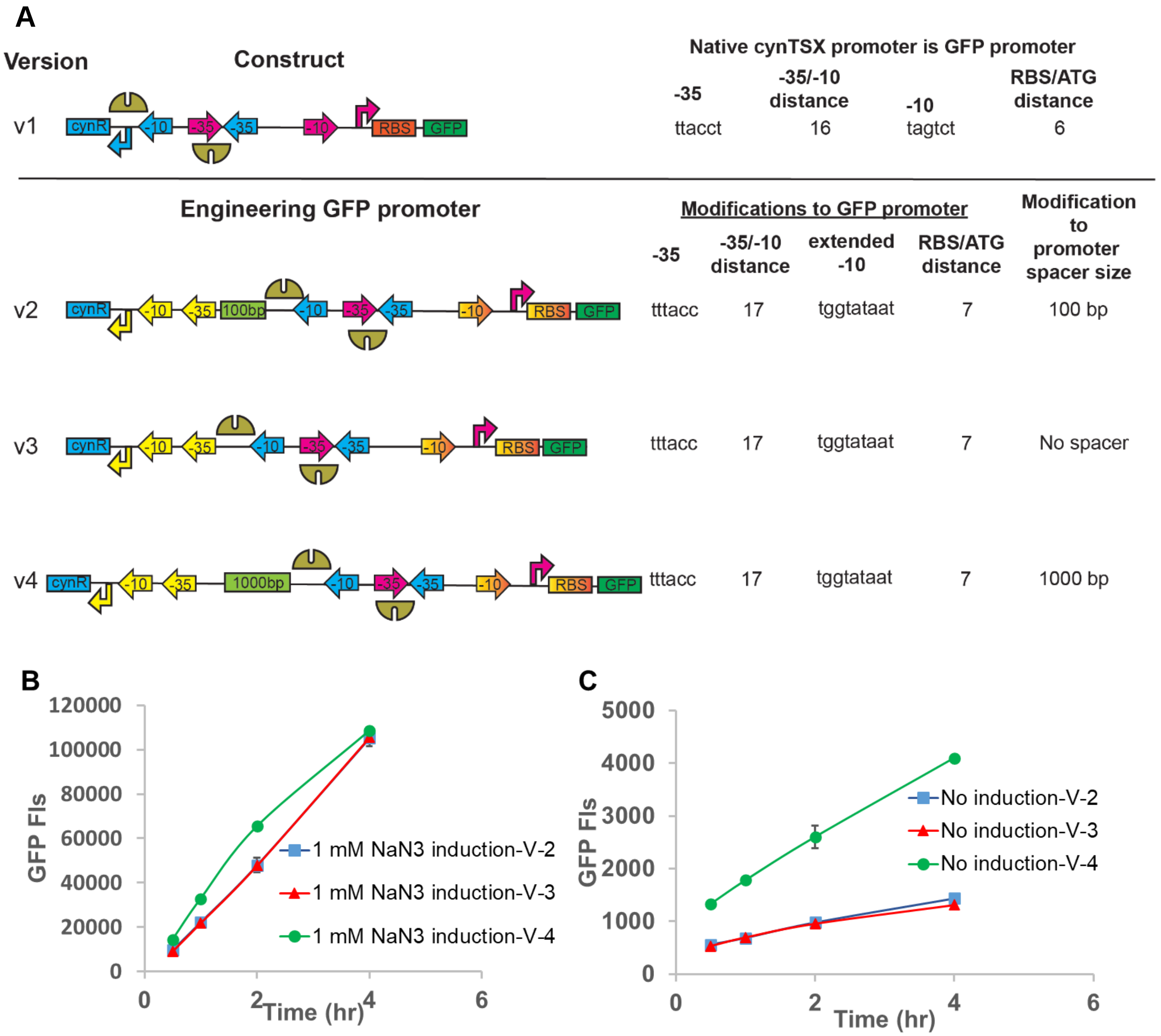

**Supplementary Figure S3.** *E. coli* cells incubated with varying concentrations of sodium azide to study the inhibitory effect of azide on cell growth. Cells were grown in LB media at 37°C for 18 hours and the OD of the cells were measured at various intermittent time points after introduction of sodium aide at time t=0. Error bars indicate one standard deviation from reported mean values from three biological replicates

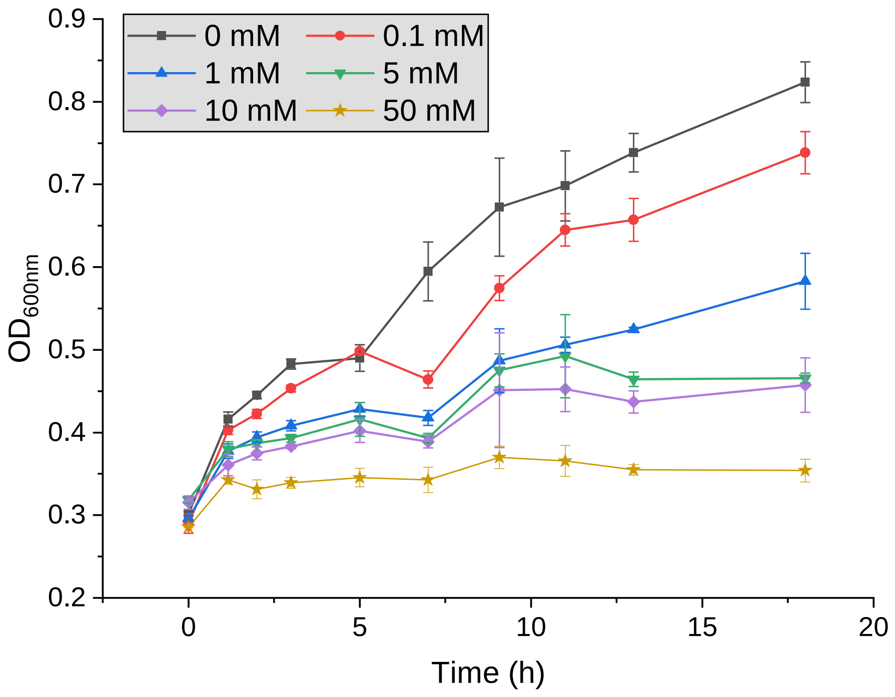

**Supplementary Figure S4.** Cell lysate GFP fluorescence from *E.coli* BW25113 cells induced using 1 mM sodium azide but at different starting cell optical density ($OD_{600}$). Cells were grown until the desired OD in LB media and induced with 1 mM sodium azide and grown then for 4 hours with measurements taken at 1 hour (grey square, left axis) and 4 hours (red square, right axis), respectively. The GFP fluorescence on the y-axis is normalized with the measured OD after induction for specified culture times. Error bars indicate one standard deviation from reported mean values from three biological replicates

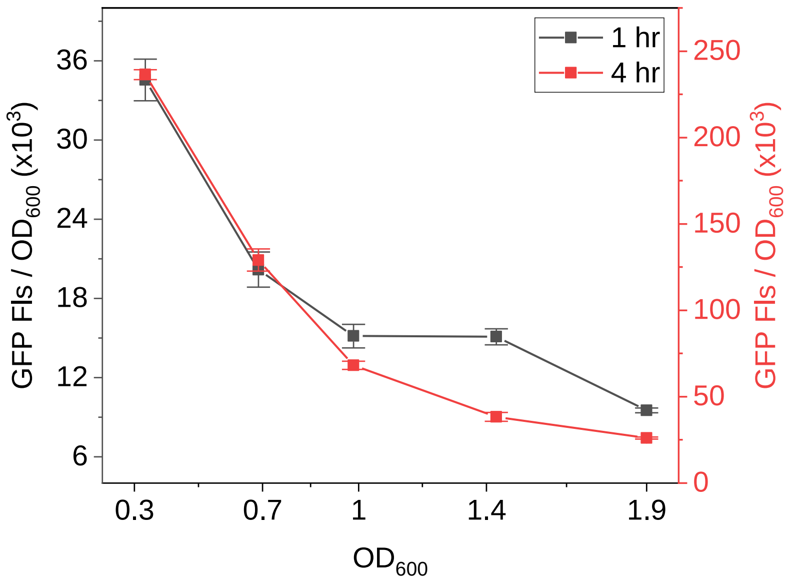

**Supplementary Figure S5. A.** Illustration of the promoter regions of pCyn-v5-GFP, pCyn-v6-GFP, pCyn-v7-GFP, pCyn-v8-GFP. **B.** Fluorescence measurements of BW25113-Wt cells with plasmids pCyn-v2-GFP, pCyn-v5-GFP, pCyn-v6-GFP, pCyn-v7-GFP incubated with 1 mM sodium azide for 2 hours. The fluorescence and OD measurements were taken after 2 hours of induction. The GFP fluorescence was normalized with the OD and plotted as bar graphs. Error bars indicate one standard deviation from reported mean values from three biological replicates

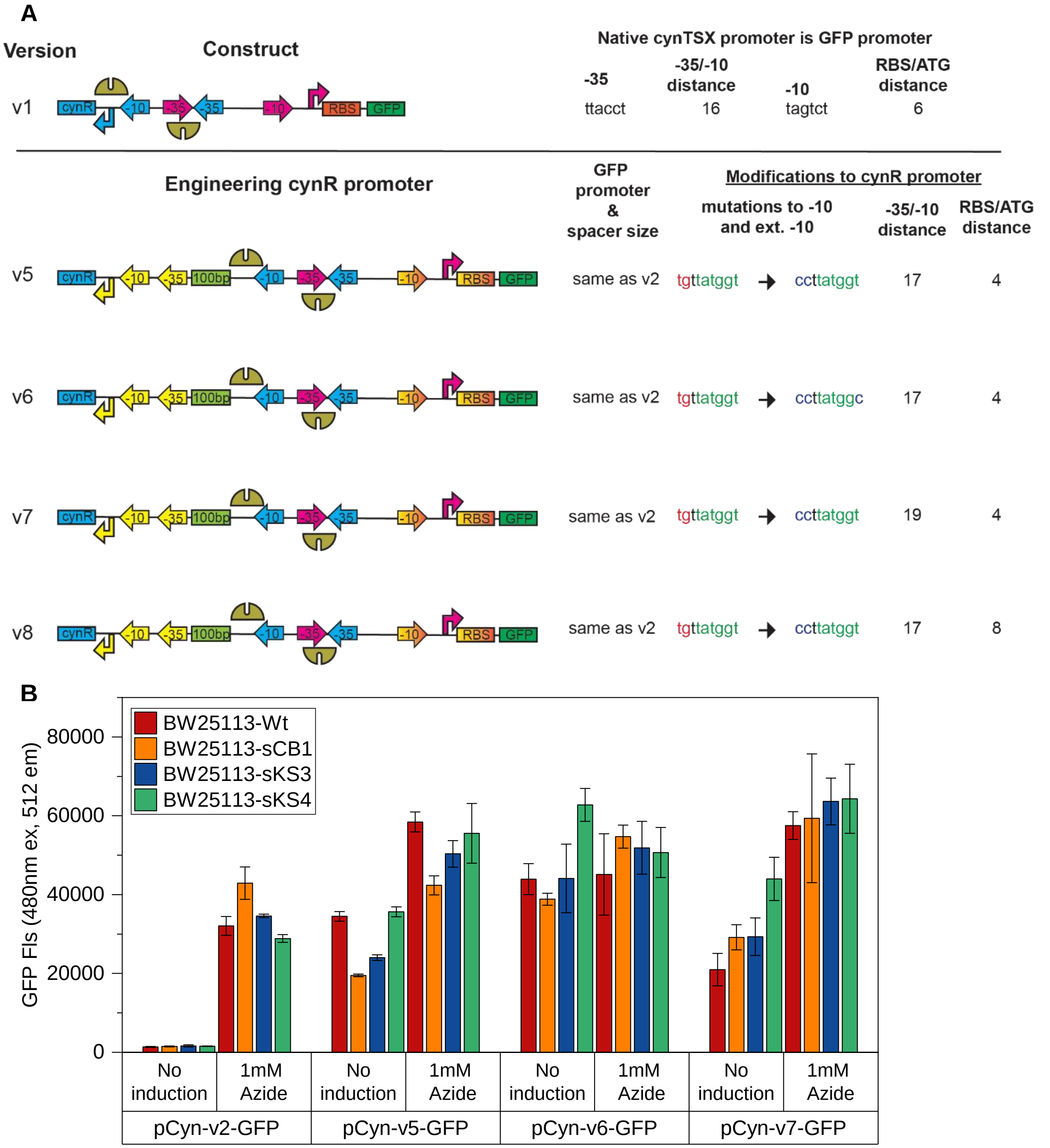

**Supplementary Figure S6**. Calibration curve for GFP protein concentration (mg/ml) and measured cell lysate fluorescence measurements.

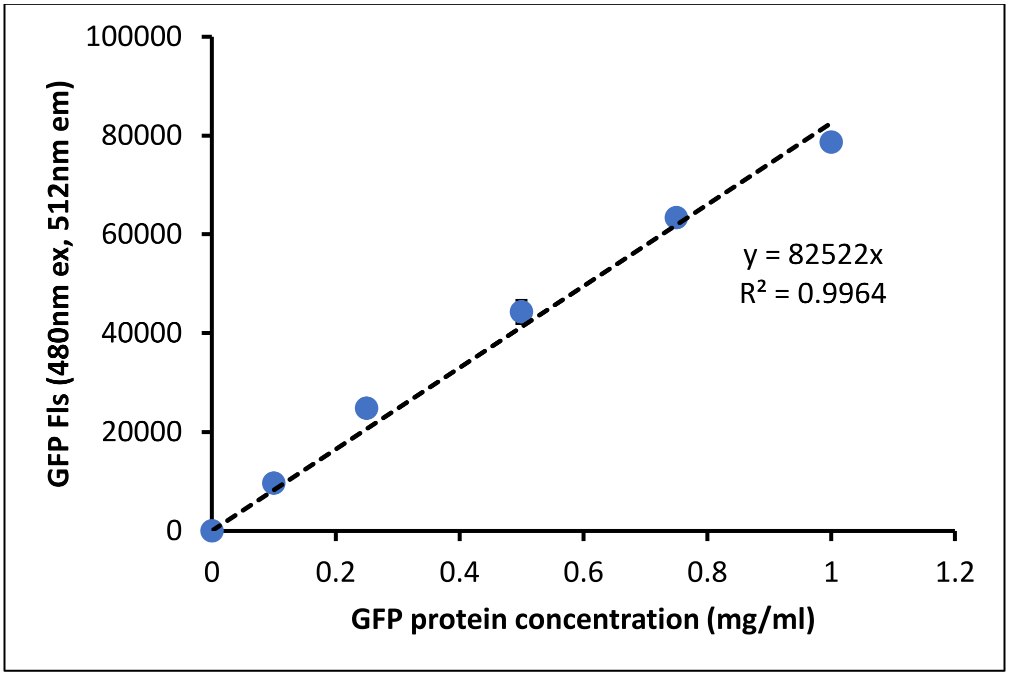

**Supplementary Text**

**>> Promoter region for pCyn-v1-GFP**

TCGCAACCTATAAGTAAATCCAATGGAACTCATCATAAATGAGACTTTTACCTTATGACAAT
CGGCGAGTAGTCTGCCTCTCATTCCAGAGACAGACAGAGGTTAACGATG

**>> Promoter region for pCyn-v2-GFP**

GGTTCCTCATTACCGTTATCATATGAACACACCATAACAAAGATGCATGCAGCTGTCTAAAT
CCCGCGGCCATGGCGGCCGGGAGCATGCGACGTCGGGCCCAATTCGCCCGATCTTAATG
AATGGCCGGAAGAGGTACGGACGCGATATGCGGGGGTGAGAGGGCAAATAGGCAGGTTC
GCCTTCGTCACGCTAGGAGGCAATTCTATAAGGATCCTCGCAACCTATAAGTAAATCCAAT
GGAACTCGTCAGAAATGAGACTTTTACCTTATGACAATCGGCTGGTATAATGCCTCTACTTC
CAGAGACAGACATAAGGAGATTACGCATG

**>> Promoter region for pCyn-v3-GFP**

GGTTCCTCATTACCGTTATCATATGAACACACCATAACAAAGATGCATGCAGCTGTCTAAAT
CCCGCGGCCATGGCGGCCGGGAGCATGCGACGTCGGGCCCAATTCGGGATCCTCGCAAC
CTATAAGTAAATCCAATGGAACTCGTCAGAAATGAGACTTTTACCTTATGACAATCGGCTGG
TATAATGCCTCTACTTCCAGAGACAGACATAAGGAGATTACGCATG

**>> Promoter region for pCyn-v4-GFP**

GGTTCCTCATTACCGTTATCATATGAACACACCATAACAAAGATGCATGCAGCTGTCTAAAT
CCCGCGGCCATGGCGGCCGGGAGCATGCGACGTCGGGCCCAATTCGCCCGATCTTAATG
AATGGCCGGAAGAGGTACGGACGCGATATGCGGGGGTGAGAGGGCAAATAGGCAGGTTC
GCCTTCGTCACGCTAGGAGGCAATTCTATAAGATTGGACCGTACGCATGTCAAACTGCTGG
CGAACCGCGATTCCACGACCGGTGCACGATTTAACTACGCCGACGTGACGACATTCCTGC
TAATGCCTCGCCCGCCGGACCGCCCTCGTGATGGGGTAGCTGGGCATGACCTTGTGACAT
ATAACGAGAGTCTACTTGTTTAATCATCTCACGGCGAAAGTCGGGGGGACAGCAGCCGCT
GCAGACATTATACCGCAACTACACCAAGCTGAGATAACTCCGTAGTTGACTACGCATCCCT
CTAGGCCTTACTTAACCGGATACAGTGACTTTGACAGGTTTGTGGGCTACAGCAATCACTT
GCATAGCTGCGTATGGAGGAAGCAACTCTTGGGTGTTAGTATGTTGACCCCTGTATTAGGG
ATGCGGGTAGTAGATGTGGGCAGAGACACCCAGGTCAAGTACACGACCCTCTCGTAGGAG
GTGTTCCAGATCACCATACCACCATACCATTCGAGCATGGCACTATGTACGCTGTCCCCAT
TCTGGTAGTCATCATCCCTATCACGGTTTCGAGTGACTGGTGACGGATATCCCCCACGAAT
GGAGATCTTATTCACAGTCGGTCACATTGGAGTGCTCCTTGACTAATCAGCTTGGCCAGGT
CTGTTGGGCCTCCGTGCCCCGAGTTTCGGCGCTGTGCTGCCGAGAGTCGGCCATTGTCAT
TGGGGCCTCACTTGTGGATACCCCGACCTATTTTGACGGGACCACTCGCGGTAGTCGTTG
GGCTTATGCACCGTGAAGTCCTCCGCCGGCCTCCCCCCTACAAAAGATGATAAGCTCCGG
CAAGCAATATTGAACAACGCAAGGATCGGCGATATAAACAGAGAAACGGCTGATTACTCTT
GTTGGTGTGGTATCGCTAAACTGGGATCCTCGCAACCTATAAGTAAATCCAATGGAACTCG
TCAGAAATGAGACTTTTACCTTATGACAATCGGCTGGTATAATGCCTCTACTTCCAGAGACA
GACATAAGGAGATTACGCATG

**>> Promoter region for pCyn-v5-GFP**

GGTTCCTCATTACCGTTATCATATGAACACACCATAAGGAAGATGCATGCAGCTGTCTAAAT
CCCGCGGCCATGGCGGCCGGGAGCATGCGACGTCGGGCCCAATTCGCCCGATCTTAATG

AATGGCCGGAAGAGGTACGGACGCGATATGCGGGGGTGAGAGGGCAAATAGGCAGGTTC
GCCTTCGTCACGCTAGGAGGCAATTCTATAAGGATCCTCGCAACCTATAAGTAAATCCAAT
GGAACTCGTCAGAAATGAGACTTTTACCTTATGACAATCGGCTGGTATAATGCCTCTACTTC
CAGAGACAGACATAAGGAGATTACGCATG

**>> Promoter region for pCyn-v6-GFP**

GGTTCCTCATTACCGTTATCATATGAACACGCCATAAGGAAGATGCATGCAGCTGTCTAAAT
CCCGCGGCCATGGCGGCCGGGAGCATGCGACGTCGGGCCCAATTCGCCCGATCTTAATG
AATGGCCGGAAGAGGTACGGACGCGATATGCGGGGGTGAGAGGGCAAATAGGCAGGTTC
GCCTTCGTCACGCTAGGAGGCAATTCTATAAGGATCCTCGCAACCTATAAGTAAATCCAAT
GGAACTCGTCAGAAATGAGACTTTTACCTTATGACAATCGGCTGGTATAATGCCTCTACTTC
CAGAGACAGACATAAGGAGATTACGCATG

**>> Promoter region for pCyn-v7-GFP**

GTTCCTCATTACCGTTATCATATGAACACACCATAAGGAAAAGATGCATGCAGCTGTCTAAA
TCCCGCGGCCATGGCGGCCGGGAGCATGCGACGTCGGGCCCAATTCGCCCGATCTTAAT
GAATGGCCGGAAGAGGTACGGACGCGATATGCGGGGGTGAGAGGGCAAATAGGCAGGTT
CGCCTTCGTCACGCTAGGAGGCAATTCTATAAGGATCCTCGCAACCTATAAGTAAATCCAA
TGGAACTCGTCAGAAATGAGACTTTTACCTTATGACAATCGGCTGGTATAATGCCTCTACTT
CCAGAGACAGACATAAGGAGATTACGCATG

**>> Promoter region for pCyn-v8-GFP**

GGAGAGTTCCTCATTACCGTTATCATATGAACACACCATAAGGAAGATGCATGCAGCTGTC
TAAATCCCGCGGCCATGGCGGCCGGGAGCATGCGACGTCGGGCCCAATTCGCCCGATCT
TAATGAATGGCCGGAAGAGGTACGGACGCGATATGCGGGGGTGAGAGGGCAAATAGGCA
GGTTCGCCTTCGTCACGCTAGGAGGCAATTCTATAAGGATCCTCGCAACCTATAAGTAAAT
CCAATGGAACTCGTCAGAAATGAGACTTTTACCTTATGACAATCGGCTGGTATAATGCCTCT
ACTTCCAGAGACAGACATAAGGAGATTACGCATG

**Supplementary Table S1: Genotype of the bacterial knockout strains generated in this study.**

| Strain name | Genotype |
|---|---|
| BW25113-wt | Wild type |
| BW25113-sCB1 | BW25113 *cynS*::FRT |
| BW25113-sKS3 | BW25113 *cynX*::FRT |
| BW25113-sKS4 | BW25113 *cynR,cynSTX*::FRT |

**Supplementary Table S2:**

**A. Primers used for performing Sequence and ligation independent cloning (SLIC) and site directed mutagenesis (SDM)**

| Primer name | Sequence |
|---|---|
| k1 | TGATAAGCGTAGCGCATCAGGCAATTCCAGCCGCAGACCTGTGTCAGCGGGT GTAGGCTGGAGCTGCTTCGAAG |
| k2 | TCTACATTAGCCGCATCCGGCATGAACAAAGCGCAGGAACAAGCGTCGCACA TATGAATATCCTCCTTAGTTCC |
| p1 | AACCAGGATAATGATACAGATTAAATCAGAACGCAGAAG |
| p2 | TTGTAATCGATAACGTAAATGCATGCCGCTTC |
| p3 | GCATTTACGTTATCGATTACAAACGTTGAACGAC |
| p4 | AATCTGTATCATTATCCTGGTTCTTTCCCCCA |
| p5 | CGGTGGCAGCTCCAAAGGTGAAGAACTG |
| p6 | AATGAGGAACCATGCTCTCTCGACATATC |
| p7 | CGAGAGAGCATGGTTCCTCATTACCGTTATCATATG |
| p8 | CCTTTGGAGCTGCCACCGCCATG |
| p9 | CGGGCCCAATTCGGGATCCTCGCAACCTATAAGTAAATCC |
| p10 | CGAGGATCCCGAATTGGGCCCGACGTC |
| p11 | CGCTAAACTGGGATCCTCGCAACCTATAAGT |
| p12 | GTACGGTCCAATCTTATAGAATTGCCTCCTAGCGTG |
| p13 | CAATTCTATAAGATTGGACCGTACGCATGTC |
| p14 | GCGAGGATCCCAGTTTAGCGATACCACACCA |
| p15 | ATGAACACACCATAAGGAAGATGCATGCAGCT |
| p16 | AGCTGCATGCATCTTCCTTATGGTGTGTTCAT |
| p17 | GTTATCATATGAACACGCCATAAGGAAGATGCA |
| p18 | TGCATCTTCCTTATGGCGTGTTCATATGATAAC |
| p19 | CATATGAACACACCATAAGGAAAAGATGCATGCAGCTGTC |
| p20 | GACAGCTGCATGCATCTTTTCCTTATGGTGTGTTCATATG |
| p21 | AGAGTTCCTCATTACCGTTATCATATG |
| p22 | CTCTCCATGCTCTCTCGACATATC |

**B. Gene block fragments used for cloning and generating plasmid constructs**

| DNA fragment name | Sequence |
|---|---|
| gblock1 | ATCGATTACAAACGTTGAACGACTGGGTTACAGCGAGCTTAGTTTATG CCGGATGCGGCGTGAACGCCTTATCCGGCCTACGTAGAGCACTGAAC TCGTAGGCCTGATAAGCGTAGCGCATCAGGCAATTCCAGCCGCTGAT CTGTGTCAGCGGCTACCGTGATTCATTCCCGCCAACAACCGCGCATT CCTCCAACGCCATGTGCAAAAATGCCTTCGCAGCGGCTGTCTGCCAG |

| | |
|---|---|
| | CTGTAGTTTATGCCGGATGCGGCGTGAACGCCTTATCCGGCCTACGT<br>AGAGCACTGAACTCGTAGGCCTGATAAGCGTAGCGCATCAGGCAATT<br>CCAGCCGCAGACCTGTGTCAGCGGCTACCGTGATTCATTTCCGCCAA<br>CAACCGCGCATTTATCCAACGCCATGTGCAAAAATGCCTTCGCGGCG<br>GCTGTCTGCCAGCTATTTTTCCGCCGCAACAAAACCGCCGTTCTCTCC<br>AGTAGTGGCGGGGCAAGAGAAATAGCTTTAAGCCCGTCATGTTGTGT<br>GGCAATCGCTGCTGGTAACAATGTGGAAAGGGAAGTGCGGCGAATCA<br>GCTCCAGAACCGCGCTAATTGAGTTCGCCTCAATGACCACCTGTGGA<br>TGTAGCCCCGCTTTCTCGCAGTAGTGGTCAATTTGCTCTCTGGTGGCA<br>AATTCCGCGCTGAGCAGGACCAGTTTTTCATCATGCAAGCGACTCAAC<br>GCCACCTGTTCATGGACGGCCAGCGGATGATGTTGCGCCACGACTAA<br>CGCTAAACTTTCTGTCAGTAAAGGAATTGCCTCCAGCTCCGGCGAATG<br>CACAGGCGCGAAGGCAATCCCAACGTCCAACTCGTCGCGGCAAAGC<br>ATATCCTCGATTTTCTCCTGCGACATTTCCTGTAGCTGGAGCGTGATG<br>CTGGGATAGCGCGCATAGAAATCCGCCATTAAGGGGCCGATAAAGTA<br>GCTCGTAAAGGTGGGGGTGACGGCGATACGCAGCGATCCTCGCGTC<br>AGATCGGCAACATCATGAATCGCCCGTTTACCCGCCCCCAGTTCCTG<br>TAACGCCCGGCTGGCGTACTGTCGCCAGACTTCTCCTGCATCAGTGA<br>GACGAATCGTTCGCCCGCTACGGTCAAACAGCGGCACGCCTAAACTC<br>TCCTCTAACTGGCGAATCTGCTGGGAAAGCGCAGGTTGGGAGACGTG<br>CAACGCACTGGCGGCACGGGTGAAGCTGCCATGTTCAGCCACGGCA<br>AGAAAATAATTGATATGTCGAGAGAGCATTCGCAACCTATAAGTAAAT<br>CCAATGGAACTCATCATAAATGAGACTTTTACCTTATGACAATCGGCG<br>AGTAGTCTGCCTCTCATTCCAGAGACAGACAGAGGTTAACGATG |
| gblock2 | GGTTCCTCATTACCGTTATCATATGAACACACCATAACAAAGATGCAT<br>GCAGCTGTCTAAATCCCGCGGCCATGGCGGCCGGGAGCATGCGACG<br>TCGGGCCCAATTCGCCCGATCTTAATGAATGGCCGGAAGAGGTACGG<br>ACGCGATATGCGGGGGTGAGAGGGCAAATAGGCAGGTTCGCCTTCG<br>TCACGCTAGGAGGCAATTCTATAAGGATCCTCGCAACCTATAAGTAAA<br>TCCAATGGAACTCGTCAGAAATGAGACTTTTACCTTATGACAATCGGC<br>TGGTATAATGCCTCTACTTCCAGAGACAGACATAAGGAGATTACGCAT<br>GCATCACCATCATCACCATCACCATGGCGGTGGCAGC |
| gblock3 | GATTGGACCGTACGCATGTCAAACTGCTGGCGAACCGCGATTCCACG<br>ACCGGTGCACGATTTAACTACGCCGACGTGACGACATTCCTGCTAAT<br>GCCTCGCCCGCCGGACCGCCCTCGTGATGGGGTAGCTGGGCATGAC<br>CTTGTGACATATAACGAGAGTCTACTTGTTTAATCATCTCACGGCGAA<br>AGTCGGGGGGACAGCAGCCGCTGCAGACATTATACCGCAACTACACC<br>AAGCTGAGATAACTCCGTAGTTGACTACGCATCCCTCTAGGCCTTACT<br>TAACCGGATACAGTGACTTTGACAGGTTTGTGGGCTACAGCAATCACT<br>TGCATAGCTGCGTATGGAGGAAGCAACTCTTGGGTGTTAGTATGTTGA<br>CCCCTGTATTAGGGATGCGGGTAGTAGATGTGGGCAGAGACACCCAG<br>GTCAAGTACACGACCCTCTCGTAGGAGGTGTTCCAGATCACCATACC<br>ACCATACCATTCGAGCATGGCACTATGTACGCTGTCCCCATTCTGGTA<br>GTCATCATCCCTATCACGGTTTCGAGTGACTGGTGACGGATATCCCCC<br>ACGAATGGAGATCTTATTCACAGTCGGTCACATTGGAGTGCTCCTTGA<br>CTAATCAGCTTGGCCAGGTCTGTTGGGCCTCCGTGCCCCGAGTTTCG<br>GCGCTGTGCTGCCGAGAGTCGGCCATTGTCATTGGGGCCTCACTTGT<br>GGATACCCCGACCTATTTTGACGGGACCACTCGCGGTAGTCGTTGGG<br>CTTATGCACCGTGAAGTCCTCCGCCGGCCTCCCCCCTACAAAAGATG<br>ATAAGCTCCGGCAAGCAATATTGAACAACGCAAGGATCGGCGATATAA<br>ACAGAGAAACGGCTGATTACTCTTGTTGGTGTGGTATCGCTAAACTG |

### References


1. Datsenko, K. A. & Wanner, B. L. One-step inactivation of chromosomal genes in Escherichia coli K-12 using PCR products. *Proc Natl Acad Sci U S A* **97**, 6640–6645 (2000).

2. Baba, T. *et al.* Construction of Escherichia coli K-12 in-frame, single-gene knockout mutants: the Keio collection. *Mol. Syst. Biol.* **2**, (2006).

3. Stevenson, J., Krycer, J. R., Phan, L. & Brown, A. J. A Practical Comparison of Ligation-Independent Cloning Techniques. *PLoS One* **8**, e83888 (2013).

4. Agrawal, A. *et al.* Click-chemistry enabled directed evolution of glycosynthases for bespoke glycans synthesis. *bioRxiv* 2020.03.23.001982 (2020). doi:10.1101/2020.03.23.001982
